# The effect of chemical distances, physical distances, and the presence of sexuals on aggression in *Cataglyphis* desert ants

**DOI:** 10.1101/478552

**Authors:** Shani Inbar, Eyal Privman

## Abstract

In social insects, non-nestmates interactions are typically agonistic and many factors may influence the degree of exhibited aggression. Two of these factors are the physical proximity between nests and the chemical dissimilarity between colonies’ chemical signatures. We studied a population sample of 43 colonies of *Cataglyphis niger* ants distributed along a transect of ∼4KM. This geographic distribution allowed us to examine correlations of aggression levels with physical and chemical distances. Ants were collected before mating season, when sexuals (unmated gynes and drones) could be found in nests. In our sample, colonies had either gynes or drones but never both. The presence of sexuals, therefore, was another factor we took into account in our behavioral analyses. Workers from nests with sexuals were more aggressive towards conspecifics than workers from nests where sexuals were absent. We also found those workers to be more vigorously active towards colonies with greater chemical distances, while workers from nests without sexuals were indifferent to chemical distances. We therefore concluded that ants are able to detect differences in chemical dissimilarity, but their aggression levels are mainly determined by other mechanisms. A possible additional mechanism is associative learning and long term memory of the chemical signatures of neighboring colonies. Such learning is supported by our finding that aggression is higher towards neighboring nests, which is in line with the previously reported ‘nasty neighbour’ effect in *Cataglyphis* ants. These results suggest that previous experience and learning of neural templates representing neighbors’ chemical cues is a stronger component than chemical dissimilarity in the mechanisms which determine aggression towards conspecifics in this species. We discuss possible explanations for the observed effect of the presence of sexuals on agonistic behavior and responsiveness to chemical distances.

## Introduction

### Nestmate recognition cues and long-term memory

Discriminating between group members and non-members is essential to the evolution and stability of sociality. In social insects, colonial identity is based on the perception of a multicomponent blend of lipids coating the cuticle of all nestmates, primarily long-chain hydrocarbons (Singer 1998, Lenoir et al. 1999, van Zweden and d’Ettorre 2010). According to the “Gestalt” model, the chemical cues are transferred among workers, creating a homogenous colony-specific chemical signature (Crozier and Dix 1979). By matching this chemical label with a neural template of the colony odour, workers are able to discriminate between nestmates and non-nestmates (Holldobler and Michener 1980, Crozier and Pamilo 1996, Vander Meer and Morel 1998, d’Ettorre and Lenoir 2010). Non-nestmates conspecific interactions are typically agonistic (Carlin and Hölldobler 1986, Soroker et al. 1994, Soroker et al. 1995a, Soroker et al. 1995b, Lahav et al. 1999, Wagner et al. 2000). Aggressive behavior between non-nestmates depends on many known factors such as genetic relatedness (Stuart and Herbers 2000, van Zweden and d’Ettorre 2010), environmental cues like nest material, climate, and queen odors (Jutsum et al. 1979, Carlin and Hölldobler 1986, Crosland 1989, Chen and Nonacs 2000), diet (Silverman and Liang 2001, Buczkowski et al. 2005, Grover et al. 2007), worker caste, task, and age (Nowbahari and Lenoir 1989, Wagner et al. 1998) etc. Two additional factors affecting levels of aggression are the degree of chemical dissimilarity between colonies (Richard et al. 2007, Martin et al. 2012) and past experience and familiarization (Errard and Hefetz 1997, Newey et al. 2010, Van Wilgenburg et al. 2010, Esponda and Gordon 2015).

The literature discusses extensively the evidence for associative learning and long-term memory in nestmate recognition. Some studies suggested an imprinting-like prenatal learning of the chemosensory stimuli (Fielde 1903, Isingrini et al. 1985, Jaisson 1991, Caubet et al. 1992, Signorotti et al. 2014). However, since colony odour is dynamic and changes over time (Vander Meer et al. 1989, Provost et al. 1993, Lahav et al. 2001, Newey et al. 2009, van Zweden et al. 2009), other studies suggested that recognition depends on plasticity and continuous updating of the template (Isingrini et al. 1985, Carlin et al. 1987, Fénéron and Jaisson 1995, Liu et al. 1998). Early learning and ongoing updating of templates, however, are not mutually exclusive mechanisms and as (Bos and d’Ettorre 2012) suggested, the template could be memorized early in life, and then maintained, reinforced or updated.

In the desert ant *Cataglyphis niger*, our model species, ants are less aggressive towards familiar conspecifics (Nowbahari 2007, Foubert and Nowbahari 2008). Such evidence supports the continuous learning hypothesis in this species. In the present study, we provide further evidence for associative learning and show that additional factors, beyond chemical similarity, play a role in eliciting aggressive behaviour towards conspecifics. We examined two of those factors; the physical proximity between colonies and the reproductive phase of the colony.

### The nasty-neighbor effect

Studies suggest that ants can discern whether a conspecific they encounter comes from a neighboring nest or a distant one, and that this affects the displayed level of aggression. In *C. niger*, workers tend to be more aggressive towards conspecifics from neighboring nests (up to 50m away), a phenomenon known as the ‘nasty neighbor’ effect (Saar et al. 2014). This phenomenon was also reported for other *Cataglyphis* species (Knaden and Wehner 2003), as well as other ant genera (Sanada-Morimura et al. 2003, Boulay et al. 2007, Thomas et al. 2007, Wilgenburg 2007, Newey et al. 2010, Roux et al. 2013). In some species, the opposite effect was reported, a phenomenon known as the ‘dear neighbor’ effect (Langen et al. 2000). Other studies, such as the study of *Formica exsecta* by (Martin et al. 2012) did not find an effect of physical distance on aggression.

### The reproductive phase of the colony

Another factor that may influence aggressive behaviour, as we will suggest in this study, is the reproductive phase of the colony and the presence of sexuals, i.e., virgin queens and males (gynes and drones). During the reproductive season, colonies may rear both gynes and drones but some ant populations exhibit a split sex ratio, where each colony produces either gynes or drones but not both (Grafen 1986, Boomsma and Grafen 1990, Boomsma 1991, Boomsma and Grafen 1991, Chan and Bourke 1994, Pamilo and Seppä 1994, Sundström 1994, Ratnieks and Boomsma 1997, Queller and Strassmann 1998, Chapuisat and Keller 1999, Mehdiabadi et al. 2003, Bourke 2005). As (Trivers and Hare 1976) suggested, split sex ratios may be the result of a tug-of-war between queens and workers over reproductive control. In colonies with singly-mated queens, queens should favor a 1:1 ratio of gynes to males and workers would prefer a 3:1 ratio. Studies from singly-mated species revealed frequent female-biased sex ratios indicating that workers often prevail (Trivers and Hare 1976, Herbers 1984, Nonacs 1986). Conversely, the queen may thwart worker control by specializing in producing either gynes or males, so that workers are unable to manipulate the sex ratio by selectively eliminating males. Determining whether colonies have a stable specialization and produce only one sex requires a long-term study because it is possible that colonies switch between female and male production in different years (perhaps due to environmental changes such as temperature fluctuations (Aron et al. 1994)) and it is also possible that the production of gynes and males is temporally offset, so that drones are produced slightly before gynes or vice versa (Keller et al. 1996). Colonies, therefore, should be examined at different times in the reproductive season and in multiple years. We have not done so in the present study because our goal was different: to test relations between aggression levels, physical distances, and chemical distances, and to account for the presence of gynes or drones in the colony as a possible additional factor influencing these relations.

## Methods

### Ants

Ants were collected between April 24 and May 10, 2016 from colonies dug along a 4 km transect in Betzet beach on the northern Israeli coastline (from N33.05162, E35.10245 to N33.07868, E35.10705). This population was previously referred to as *Cataglyphis drusus* (Eyer et al. 2017) but our recent species delimitation study suggested that this is the same species as *C. niger*, because these populations are not differentiated by their nuclear genomic DNA (Brodetzki et al. 2019). Colonies of this population are monogyne (headed by a single queen) and polyandrous (queens are multiply mated) (Eyer et al. 2017, Brodetzki et al. 2019). Queens usually mate during the spring and sexuals (gynes and drones) can usually be found in nests in April-May. In this study, on the days when ants were collected, we observed colonies with either gynes or males (or neither of the two) present in the colony before mating season. We collected at least 30 ants from each of 50 nests, 43 of which were used in this study. In 14 colonies we found gynes to be present, in 7 colonies we found males, and in 22 colonies we found no sexuals.

### Behavioural bioassays

For 65 pairs of colonies, 4-7 different ants from one colony were subjected to dyadic aggression assays with 4-7 different ants from another colony, a total of 330 assays. All encounters were conducted in a neutral arena (9 cm diameter, lined with a filter paper that was changed after each test). Before each test, workers were acclimatized to the arena for 2 min by placing them on the arena surface covered in separate glass tubes. The test began by removing the tubes, and aggressive behavior was recorded for 2 min. Aggression was scored according to the commonly practiced method, as in (Errard and Hefetz 1997), for the following behavioral acts: 1 point - antennation; 2-mandibular threat; 3-short biting with jumping; 4-biting; 5-spraying of formic acid. The aggression index was calculated as follows:

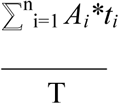

where *A*_*i*_ and *t*_*i*_ are the aggression score and duration of each of the five types of exhibited behaviors, and T is the total interaction time (120 sec).

### Cuticular hydrocarbon analysis

Cuticular hydrocarbons were measured for 4-7 workers, which were not used in the behavioral assays, from each of the 43 colonies. Whole bodies were individually immersed in hexane to extract non-polar cuticular lipids from each sample separately. Initial analysis was conducted by gas chromatography/mass spectrometry (GC/MS), using a VF-5ms capillary column, temperature-programmed from 60°C to 300°C (with 1 min initial hold) at a rate of 10°C per min, with a final hold of 15 min. Compounds were identified according to their fragmentation pattern and respective retention time compared to authentic standards. Subsequently, all samples were assayed quantitatively by flame ionization gas chromatography (GC/FID), using the above running conditions. Peak integration was performed using the program Varian Galaxie 1.9. and relative amounts of each compound were calculated. Chemical distances between colonies were determined by Mahalanobis distances calculated following a discriminant analysis.

### Data and Statistical analysis

Statistical analysis was performed using XLSTAT (https://www.xlstat.com; Addinsoft 2019;Boston, USA).

## Results

We performed a total of 330 aggression tests between non-nestmates from nests where either gynes (F), males (M), or no sexuals (N) were found: 45 FF interactions between 10 different colonies, 38 MF interactions between 8 different colonies, 27 MM interactions between 5 different colonies, 87 NF interactions between 17 different colonies, 34 NM interactions between 7 different colonies, and 99 NN interactions between 18 different colonies. We also performed 173 tests between nestmates. Physical distances between tested nests ranged between 22m and 1610m with and average distance of 387.6m. A total of 235 chemical analyses were performed on 4-7 ants from all 43 colonies. We identified 34 compounds in the long-chained hydrocarbon fraction (C25-C33), including n-alkanes, monomethyl alkanes, dimethyl alkanes and trimethyl alkanes.

### The presence of sexuals

Nestmates and non-nestmates interactions showed significant differences in aggression levels (Mann-Whitney two-tailed test: *p* < 0.0001) as expected. An overall comparison of aggression scores between N-N, N-S, and S-S (Figure 1) revealed that levels of aggression varied depending on whether workers from sexuals-producing colonies were involved. S-S interactions were most aggressive (mean score: 1.35±1), N-S interactions followed (mean score: 1.00±1.04), and N-N interaction were the least aggressive (mean score: 0.84±0.94). The S-S interactions were significantly more aggressive than both the N-S interactions and the N-N interactions (Kruskal-Wallis test followed by pairwise comparisons: The S-S vs. N-S and S-S vs. N-N comparisons both had *p* < 0.0001, whereas the N-N vs. N-S comparison had *p* = 0.17). For further subdivision by sexual types see Supplementary Figure S2 and Table S2 which show FM interactions to be the most aggressive and statistically different from other types of interactions except FF interactions.

**Figure 1:**
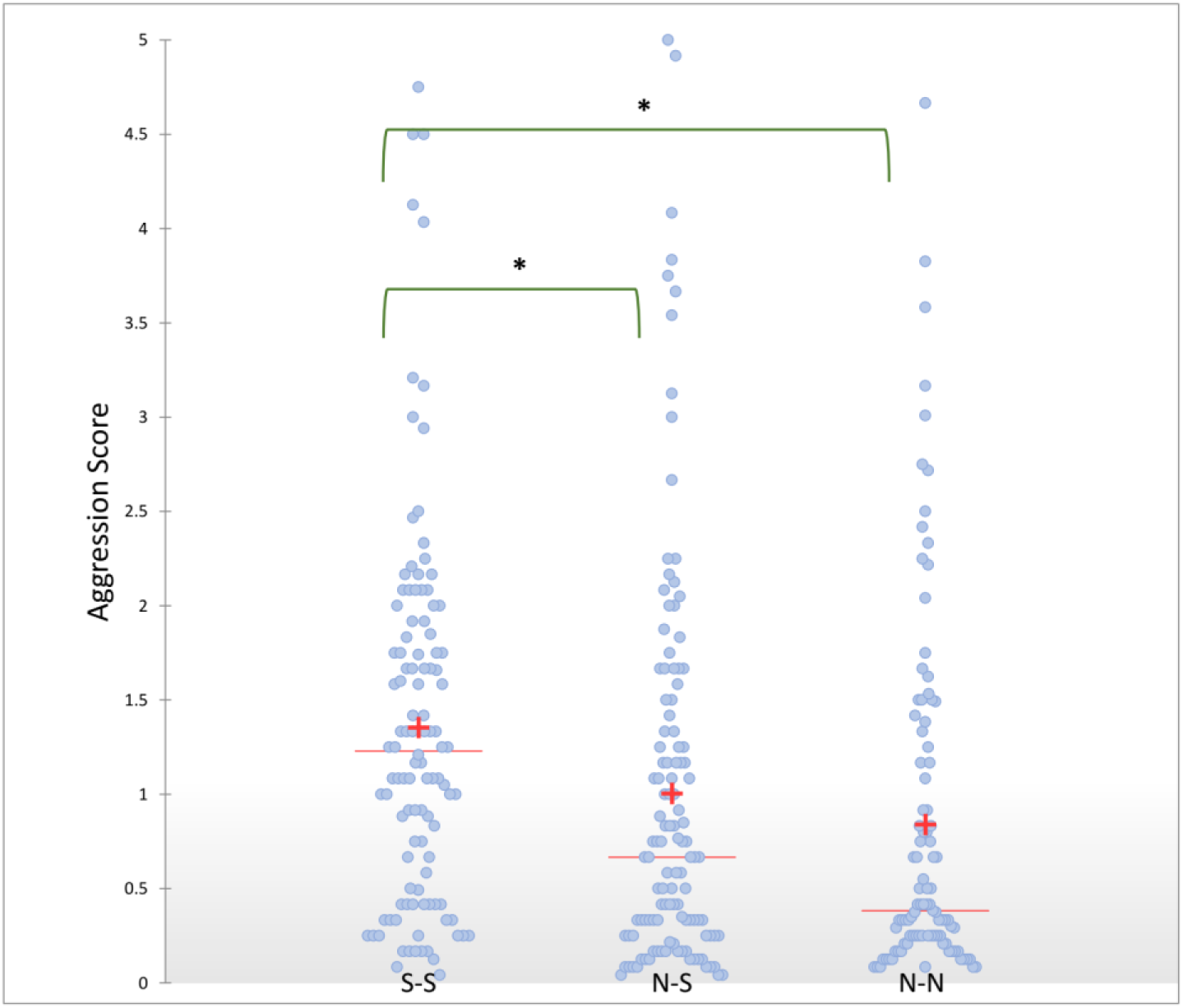
Distributions of aggression scores for three categories of interactions: between workers from nests where sexuals were present (S-S), between workers from nests where sexuals not present (N-N), and between workers from the two different categories (N-S). S-S interaction are significantly more aggressive than the other two categories.

### Chemical distances

Chemical distances between nests ranged between 13.69 and 159.52, with mean and standard deviation of 59.52±30.58. We initially found no significant correlation between chemical distances and physical distances between nests (Mahalanobis distances calculated following a discriminant analysis which can be found in Supplementary Figure S1; Spearman correlation *rho*=0.061 *p*=0.268). Interestingly, the relation between aggression and chemical distances varied between the above mentioned three categories, when aggression scores were separated based on the presence of sexuals (Figure 2): A significant positive correlation was found only in the S-S category (Spearman correlation *rho*=0.249, *p*=0.009), whereas the N-N and the N-S categories displayed a weak non-significant positive correlation (*rho*=0.096, *p*=0.345 and *rho*=0.004, *p*=0.965, respectively). Thus, when sexuals are present in the colony workers seem to respond to the chemical distance with elevated aggression, whereas workers from colonies with no sexuals do not respond to the chemical distance. Further subdivision by the type of sexuals that were found in the colony are not presented since sample became too small statistical analysis (results can be found in Supplementary Figures S3 and Table S3 Which show MM interactions to be statistically different from all other types of interactions).

**Figure 2:**
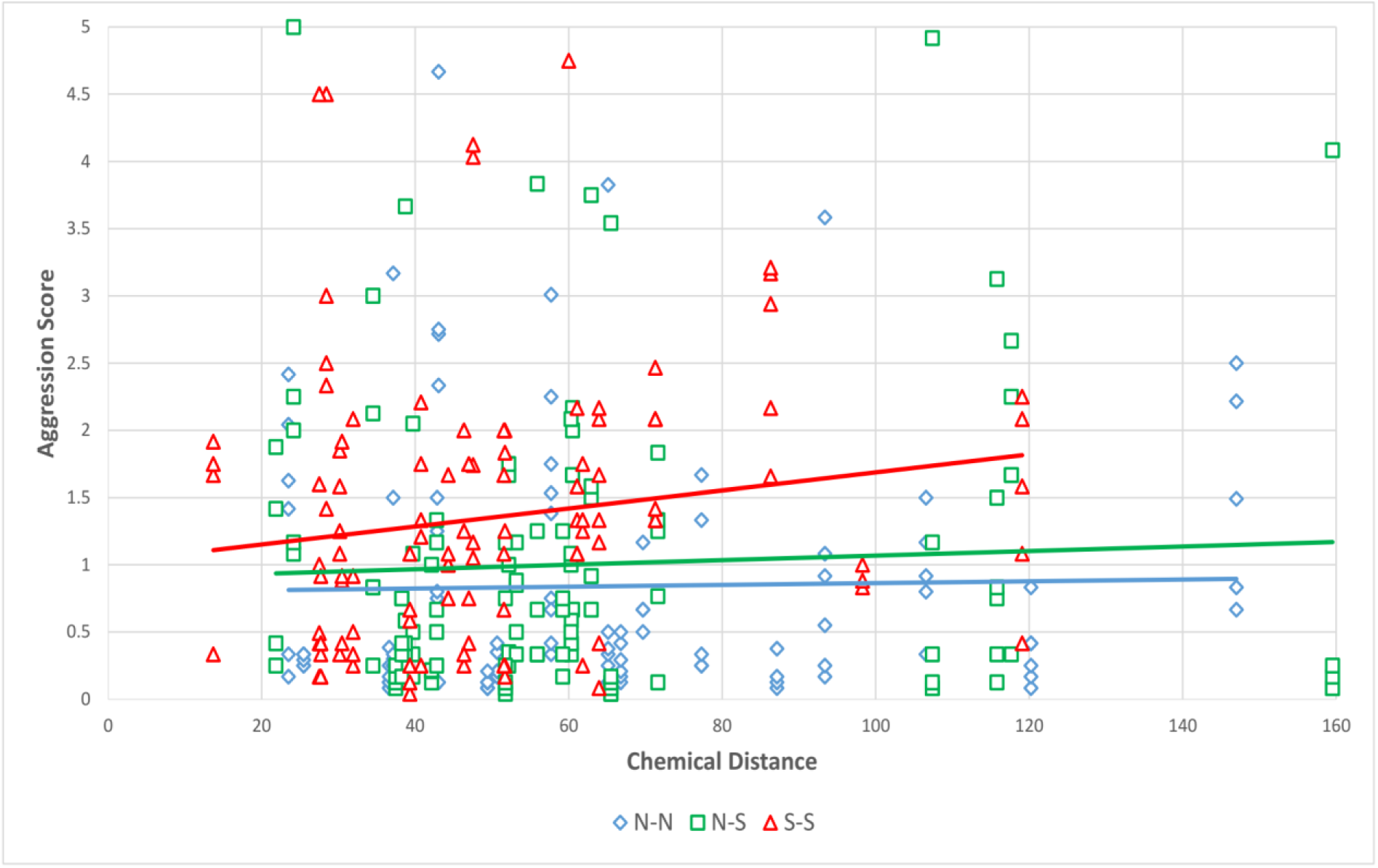
Association between aggressive behaviour and chemical distances. The three categories are defined in the legend of Figure 1. Significant positive correlation was found for the S-S interactions but not for the other categories.

### Physical distances

Next, we looked for associations of the above factors with physical distances between nests. We first plotted aggression levels as functions of physical distances (Figure 3), which exhibited a negative trend in short distances up to 50m, and then stabilized for longer distances (*p*=0.001 in Kruskal-Wallis test between two intervals, up to 50m and over 50m). This negative trend in short distances is in line with the previously reported ‘nasty neighbour’ effect (Saar et al. 2014). However, there was no linear correlation over the full range of physical distances (Spearman correlation *rho*=-0.077, *p*=0.164). Interestingly, the relation between physical distances and chemical distances showed a different picture - there was no statistically significant difference between chemical distances for the short (<50m) and long physical distances (Kruskal-Wallis *p*=0.267). However, chemical and physical distances were correlated over the full range of physical distances (Figure 4; Spearman correlation *rho=*0.248, *p*< 0.0001).

**Figure 3:**
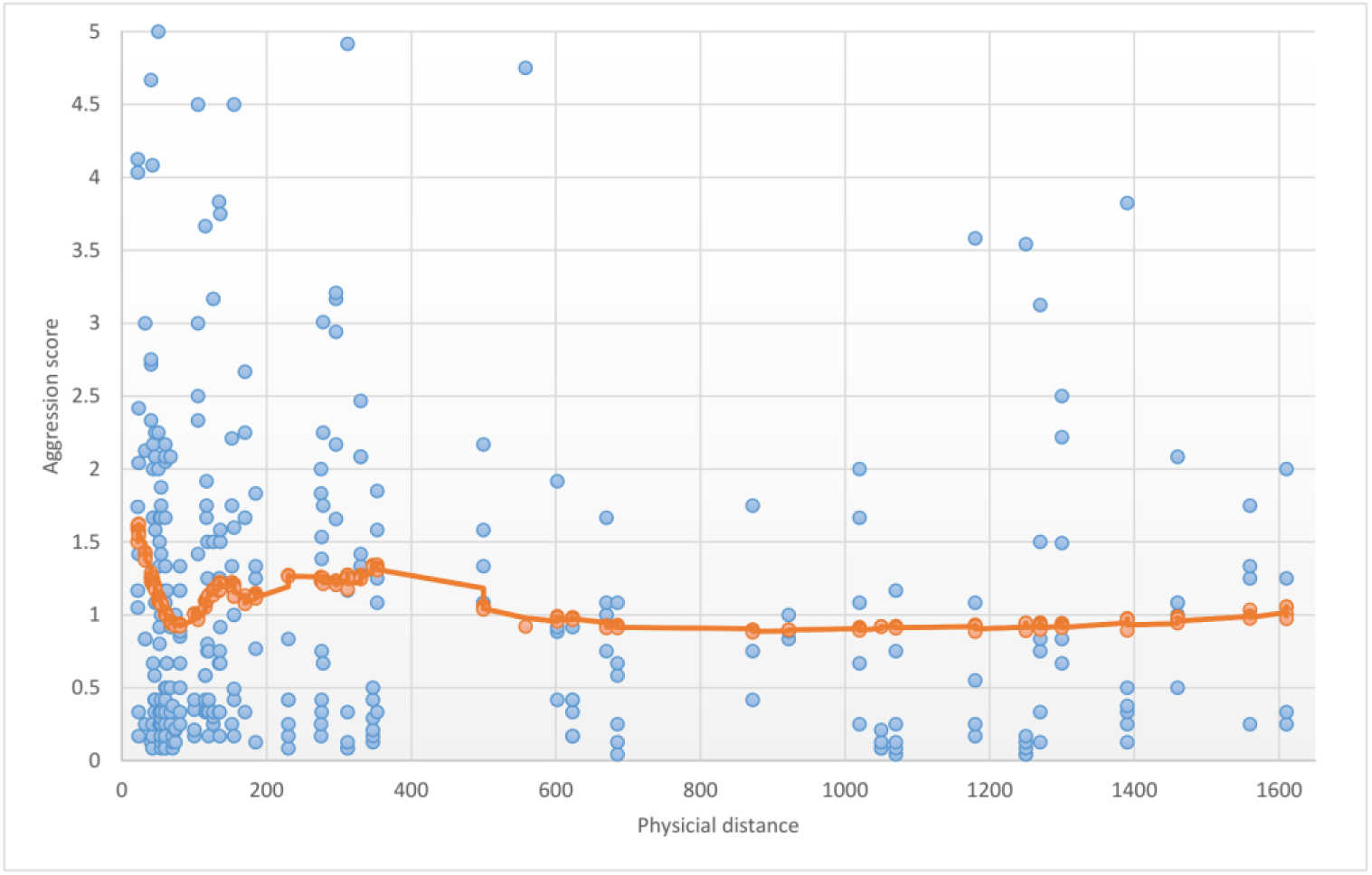
Distribution of aggression scores along the physical distances between nests. (where the encountering ants originated). Orange line indicate trend according to a LOWESS regression.

**Figure 4:**
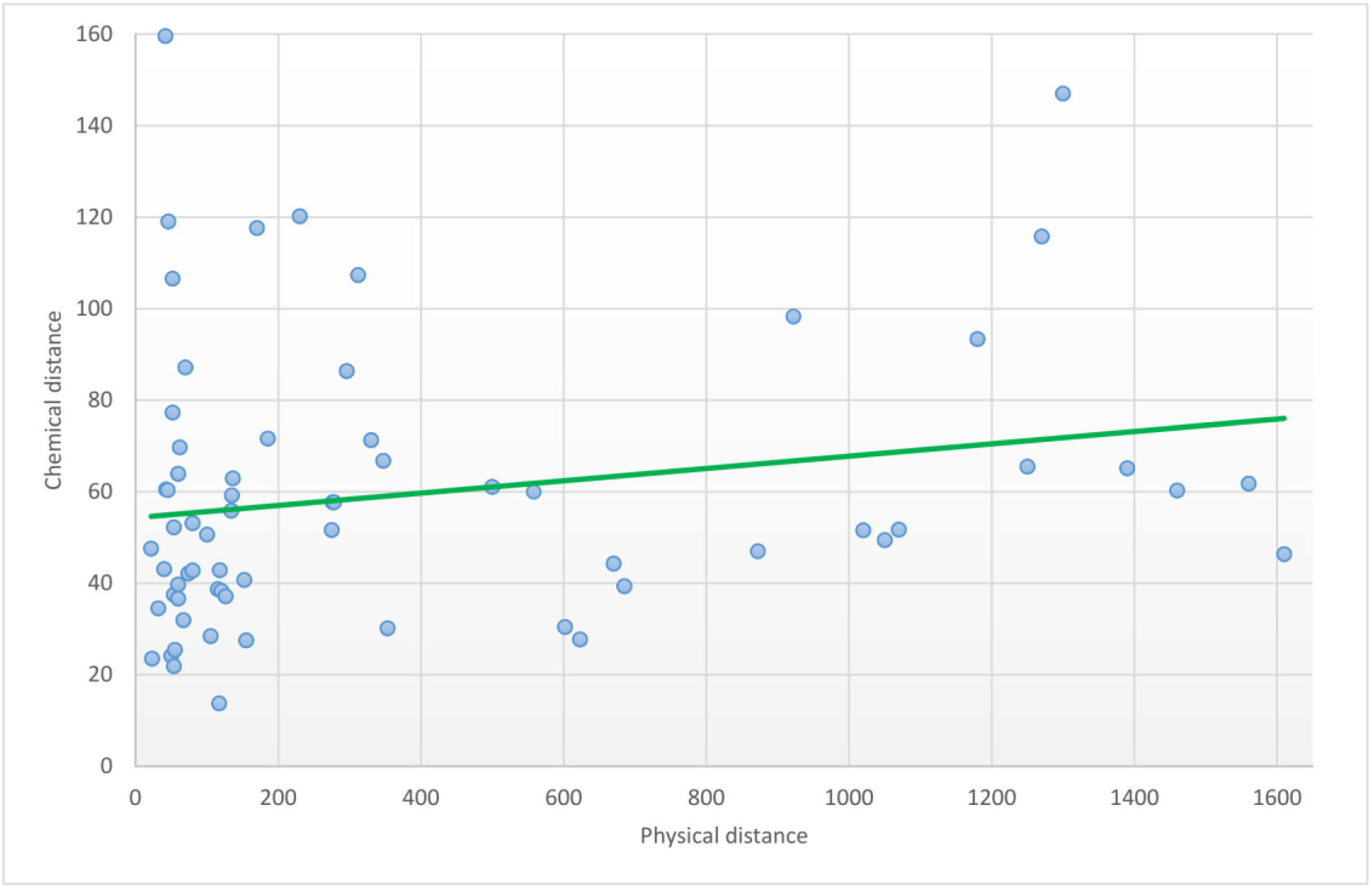
Distribution of chemical distances along the physical distances between nests. Green line indicates the linear trend.

## Discussion

In this study, we tested for effects of physical distances and chemical distances on aggression between conspecifics. Furthermore, we accounted for the presence of gynes and drones before mating seasons as an additional factor affecting these distances. We found that the level of aggression between workers from colonies where sexuals (either gynes or drones) were present was significantly elevated when compared to colonies without sexuals. We also found that when sexuals were present, aggression was correlated with the chemical distances between colonies. We interpret this result as evidence for the ants’ ability to detect chemical distances despite the fact that no such correlation was observed in ants from colonies where no sexuals were found. There are two reasons for this interpretation. Firstly, there is the inherent problem of non-significant results, which do not exclude an effect that was not detected due to limited statistical power. Secondly and more importantly, it is unlikely that in the short span during the year when sexuals are present in the colony, ants are able to detect chemical distances, and then lose that ability when sexuals are absent. We suggest that although ants have that ability throughout the year, their response to this stimulus was only detectable when their general aggressiveness was increased in the presence of sexuals. This interpretation is also in line with previous studies that found aggression to be correlated with chemical distance, at least in some species like *Formica exsecta* and *Temnothorax unifasciatus* (Foitzik et al. 2007, Martin et al. 2012).

We argue that the reproductive phase of the colony is an important intrinsic factor affecting worker response threshold to conspecific threats. The elevated aggression and response to chemical distances before mating season may be related to the high energetic toll that sexuals take and the need to protect this investment; gynes and drones are typically larger in size than the average worker and they need to develop flight muscles and store energy reserves for flight. Gynes also need to accumulate fat and protein in order to decrease, or altogether avoid, foraging during colony founding (Keller and Passera 1988, 1989). Such an energetic investment by the colony needs to be protected and thus results, we argue, in elevated worker aggression during the period between the production of sexuals and their departure for mating. This hypothesis is supported by evidence for seasonal variation in aggression levels between workers (Ichinose 1991, D’Ettorre et al. 2004, Brandt et al. 2005, Katzerke et al. 2006, Thurin and Aron 2008), which is determined by both intrinsic (e.g. worker density, relatedness within the nest) and extrinsic (e.g. temperature, availability and quality of food) (Thurin and Aron 2008) factors. Additional evidence also indicated that some ant species have periodic variation in cuticular hydrocarbon profiles (Vander Meer et al. 1989, Nielsen et al. 1999, Liu et al. 2001), which may alter the response towards non-nestmates.

When sexuals are absent, we suggest, aggression is modulated by other mechanisms and involves multiple factors. One such factor is the physical distance between nests. As reported previously in *C. niger* ants (Saar et al. 2014), we found a ‘nasty neighbour’ effect which occurs when ants respond more aggressively towards conspecifics from physically closer nests. When agonistic behaviour was examined in relation to physical distances between nests, there was a negative trend in short distances (up to 50m) which then stabilized. This elevated aggression towards neighbors may have an adaptive benefit when neighbouring nests compete over foraging areas and other resources. When conspecifics are encountered in the field, rather than forced to share a small arena as in our behavioural essay, ants usually avoid interaction altogether. The nasty neighbor effect may induce ants to attack foragers from neighboring colonies when encountered in their foraging territory. However, we found no significant difference between the chemical distance of colonies with short (up to 50m) and longer (over 50m) distances between them. There was a positive correlation between chemical and physical distances over the entire range of physical distances, but this positive trend is opposite of the negative association between aggression and physical distances (in short distances). The ‘nasty neighbour effect’, therefore, cannot be explained by the ants’ ability to detect chemical distances.

Associative learning and long term-memory, on the other hand, may explain our observations. In all likelihood, ants from neighboring nests that were used in our behavioral assays had previous experience and were familiar with each other’s colony recognition cues. We suggest that the ants learn to recognize the specific chemical profiles of their neighbors. Such associative learning could be based on the same label-template mechanism that was suggested to underlie nestmate recognition. The difference in the use of the two mechanisms, i.e., chemical dissimilarity relative to self vs. learning of the neighbor’s profile, under different social/temporal conditions could also be explained by a change in the colony’s interests. Throughout the year, when sexuals are absent, it may be in the interest of the colony to discriminate between conspecifics from close and distant nests but not as important to discern between equally close nests. Such interests would be best served by learning neighbors’ profile. Before mating season, however, other factors may influence the level of aggression. One such factor is genetic relatedness. It is possible the queens do not stray far from their mother’s nests to found their own. This would result in higher genetic relatedness among neighboring nests. If the chemical profile of the colony has a genetic basis, as studies suggest (Vander Meer and Morel 1998) then, neighboring nests would tend to have similar profiles. Indeed, this hypothesis is supported by our finding of a positive correlation between chemical and physical distances. In such a case, chemical dissimilarity and physical distance are ineffective mechanisms. We argue that these two mechanisms - discerning between non-kin by means of chemical dissimilarity and by means of associative learning - are not mutually exclusive. It seems that these two independent mechanisms are used to different extents in different contexts and depend on the internal state of the colony, in this study its reproductive phase.

## Acknowledgments

We thank Abraham Hefetz for introducing us to these wonderful ants and for assistance with the chemical analysis. E.P. was supported by the Israel Science Foundation Grants no. 646/15, 2140/15, and 2155/15 and US-Israel Binational Science Foundation Grant no. 2013408.

