## Supplemental material for "The effect of chemical distances, physical distances, and the presence of sexuals on aggression in *Cataglyphis* desert ants"

**Supplementary materials**

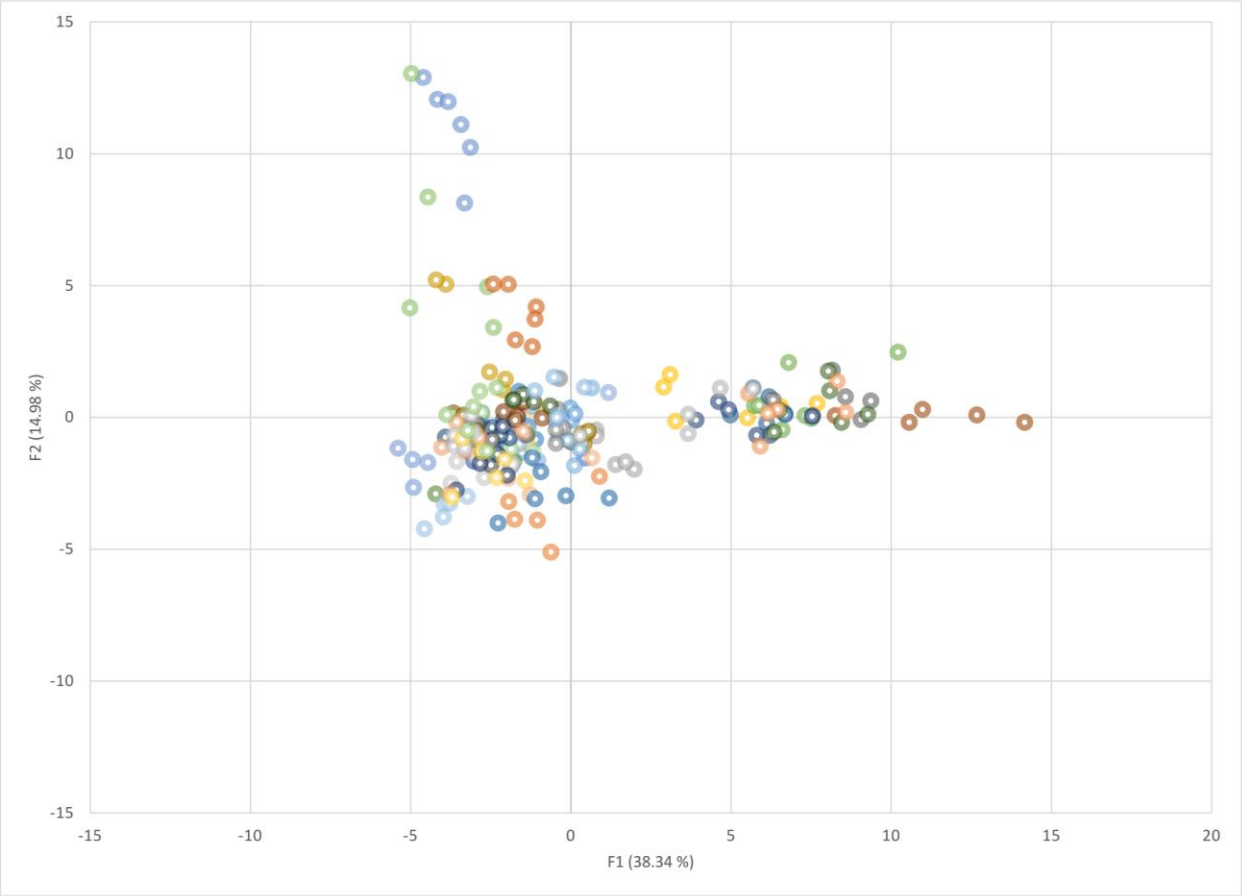


Supplementary Figure S1: Discriminant analysis (DA) of the chemical signature of all colonies examined in this study. Wilks' Lambda test (Rao's approximation) indicate a statistical difference (Lambda= 0.000, p-value<0.0001).

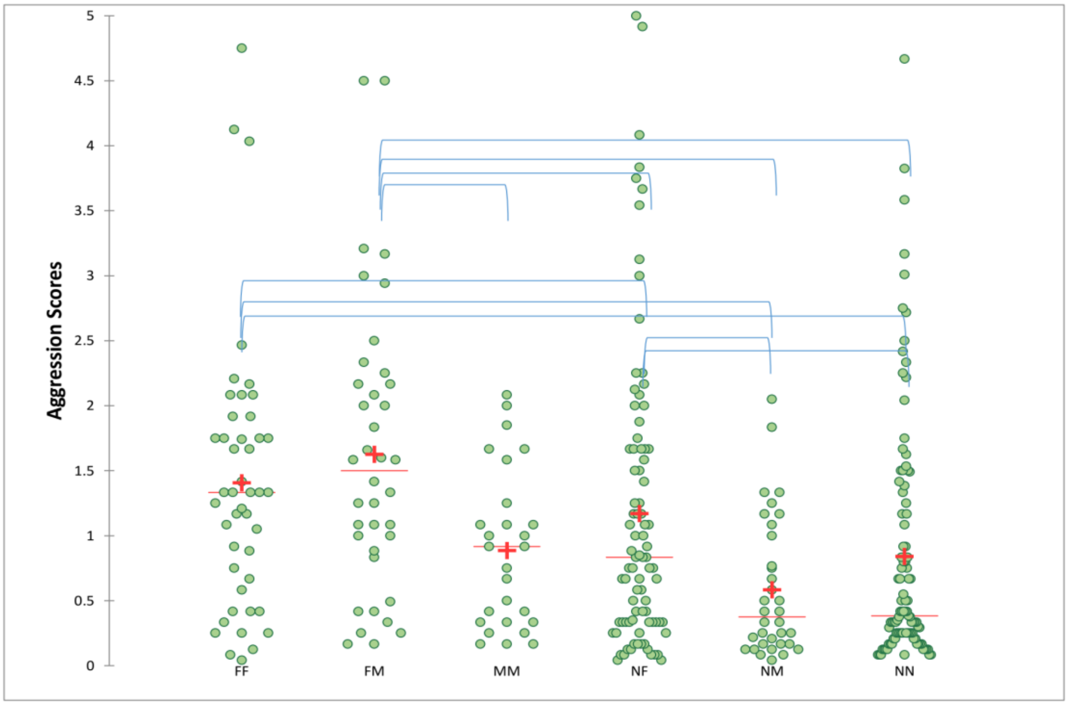


Supplementary Figure S2: Comparison of aggression scores between workers from female producing colonies (F), male producing colonies (M) or colonies where no sexulas were found (N). Blue brackets indicate pairs of statistically different groups. P-values are given in Supplementary Table S2.

Supplementary Table S2: P-values for each comparison according to Kruskal-Wallis test followed by a pairwise comparison

| | FF | FM | MM | NF | NM | NN |
|---|---|---|---|---|---|---|
| FF | 1 | 0.456 | 0.070 | 0.046 | < 0.0001 | < 0.0001 |
| FM | 0.456 | 1 | 0.016 | 0.006 | < 0.0001 | < 0.0001 |
| MM | 0.070 | 0.016 | 1 | 0.735 | 0.056 | 0.213 |
| NF | 0.046 | 0.006 | 0.735 | 1 | 0.005 | 0.019 |
| NM | < 0.0001 | < 0.0001 | 0.056 | 0.005 | 1 | 0.263 |
| NN | < 0.0001 | < 0.0001 | 0.213 | 0.019 | 0.263 | 1 |

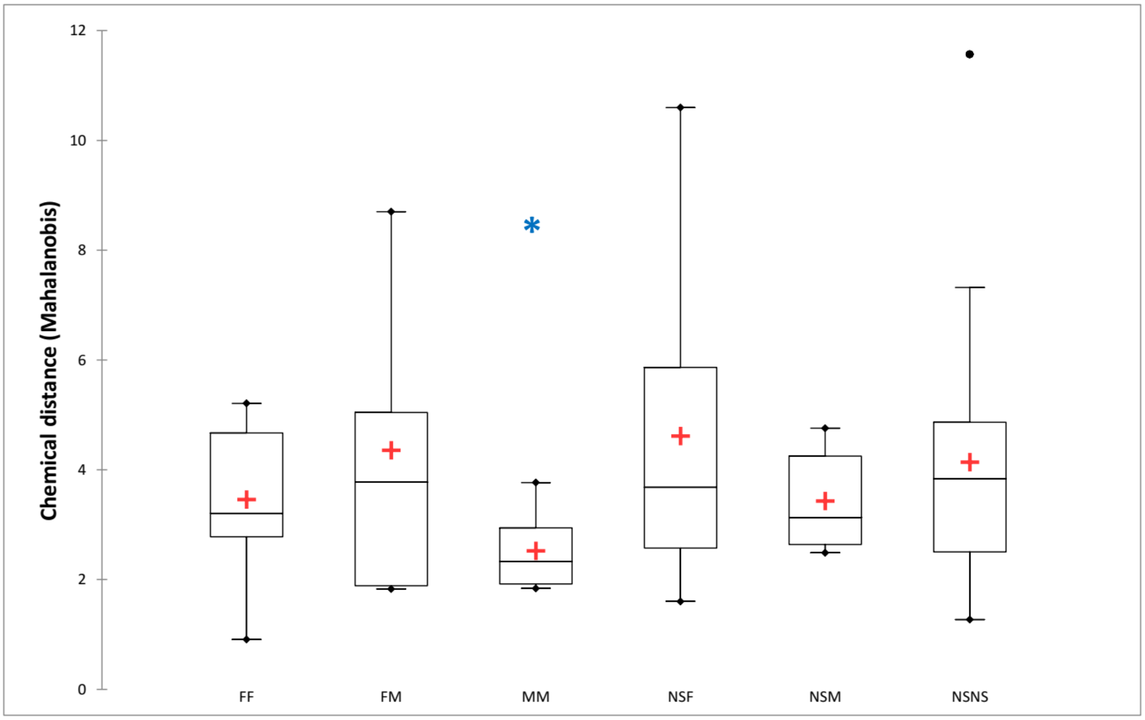


Supplementary Figure S3: Comparison of chemical distances between workers from female producing colonies (F), male producing colonies (M) or colonies where no sexulas were found (N). Blue asterisk indicate the statistically different pair. *P-values* are given in Supplementary Table S3.

Supplementary Table S3: *P-values* for each comparison according to Kruskal-Wallis test followed by a pairwise comparison

| | **FF** | **FM** | **MM** | **NF** | **NM** | **NN** |
|---|---|---|---|---|---|---|
| **FF** | 1 | 0.362 | 0.001 | 0.217 | 0.870 | 0.518 |
| **FM** | 0.362 | 1 | < 0.0001 | 0.894 | 0.314 | 0.658 |
| **MM** | 0.001 | < 0.0001 | 1 | < 0.0001 | 0.003 | < 0.0001 |
| **NF** | 0.217 | 0.894 | < 0.0001 | 1 | 0.192 | 0.452 |
| **NM** | 0.870 | 0.314 | 0.003 | 0.192 | 1 | 0.440 |
| **NN** | 0.518 | 0.658 | < 0.0001 | 0.452 | 0.440 | 1 |

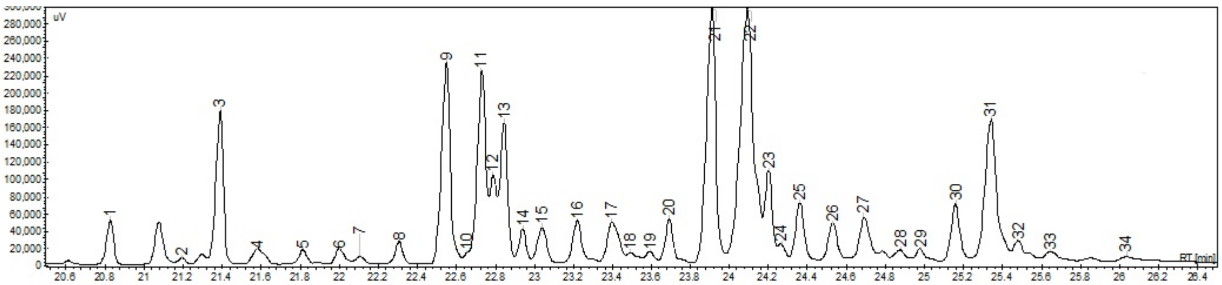


Supplementary Figure S4: Chromatograms of CHCs from total body extracts. Only long-chained CHC are shown.

Supplementary Table S4: 34 compounds that were identified in the long-chained hydrocarbon fraction

| | | | | | | | |
|---|---|---|---|---|---|---|---|
| 1 | c25 | 11 | 1,15 dime c27 | 21 | 11+13 me c29 | 31 | 11,15 dime c31 |
| 2 | 5me c25 | 12 | 7,11 dime c27 | 22 | 11,15 dime c29 | 32 | 7,11,15 trime |
| 3 | 3me c25 | 13 | 3me c27 | 23 | 3me c29 | c31 | |
| 4 | c26 | 14 | 7,11,15 trime c27 | 24 | 7,11,15 trime | 33 | c32 |
| 5 | 10me c26 | 15 | c28 | c29 | | 34 | c33 |
| 6 | 4me c26 | 16 | 12me c28 | 25 | c30 | | |
| 7 | 4,12 dime c26 | 17 | 4me +8+12 dime c28 | 26 | 14 me c30 | | |
| 8 | c27 | 18 | 2me c28 | 27 | 4 me c30 | | |
| 9 | 11+13 me c27 | 19 | 4,12 dime c28 | 28 | 2 me c30 | | |
| 10 | 5me c27 | 20 | c29 | 29 | c31 | | |
| | | | | 30 | 13+15 me c31 | | |
